# Adaptive disorder as the hallmark of nanobodies antigen-binding loops

**DOI:** 10.1101/2025.10.10.681624

**Authors:** Davide Bagordo, Gauthier Trèves, Mariangela Santorsola, Giorgio Colombo, Francesco Lescai

**Author notes:** these authors have contributed equally to the work (Davide Bagordo, Gauthier Trèves).

## Abstract

Nanobodies are antigen-binding proteins of great interest as diagnostics and therapeutics. Accurate and fast characterization of their complementarity-determining regions (CDRs) is crucial to uncover the principles guiding their design. Yet, this task remains challenging, as random recombination and somatic mutations generate highly diverse CDR sequences that escape motif-based or structure-prediction approaches currently used to identify them.

To overcome this hurdle, we employed two independent strategies that converged on the same conclusion. At the sequence level, we developed a deep learning model to identify nanobody CDRs directly from the primary sequence. At the structural level, we applied an energy decomposition method, revealing CDRs as residues highly uncoupled to the rest of the fold. Explainability analyses showed the network captured intrinsic CDR properties, which notably aligned with these energy values. CDRs emerge as fuzzy regions capable of adopting diverse conformational ensembles, from which a preferred state is selected upon antigen binding.

This finding supports a model where chaos in both sequence and structure appears adaptive and disorder emerges as the hallmark of nanobody CDRs. This work aims to advance the definition of rules for the design of antigen binding regions, paving the way for the next-generation immune diagnostics and therapeutics.

## Introduction

Protein–protein interactions (PPIs) are fundamental determinants of virtually all biological processes. Within the immune system, precise molecular recognition between host proteins and pathogenic antigens initiates cascades of biochemical events that ultimately drive the clearance of invading agents. Deciphering the sequence and structural principles that underlie these interactions is therefore central to understanding the mechanisms of immune defence.

From a conceptual standpoint, insights into the evolutionary dynamics of sequences and the molecular origins of immunological recognition can illuminate why particular biomolecules are selected as effective mediators and define the structural features that govern their binding interfaces. From a translational perspective, such knowledge provides a framework for establishing molecular design rules, thereby advancing the development of next-generation biologics—a field that has gained remarkable momentum in recent years.

In this context, Nanobodies (Nbs) have been attracting growing interest. Nbs are a class of antigen-binding fragments naturally present in camelid and shark serum^1^. They are also called V_H_H antibodies, since they are constituted only of the antibody heavy chain. They show the same architecture as antibody V_H_ domains, with four conserved framework regions (FR 1/2/3/4) around three hypervariable loops, called complementarity-determining regions (CDR 1/2/3). CDRs, along with a small number of framework residues, form the paratope region responsible for the interaction with the antigen epitope. Despite the involvement of all three loops, the main actor in epitope recognition and binding is CDR3. This region is much longer than the corresponding one in conventional antibodies, and it often adopts long fingerlike conformations able to interact inside cavities formed on antigens. Consequently, epitopes targeted by Nbs are often different from those targeted by conventional antibodies^2^.

Moreover, the smaller size of nanobodies as well as the different amino acidic composition results in numerous advantages: a higher stability and water solubility, and a stronger binding affinity for the antigen^3^. These unique features make Nbs very attractive as next-generation biodrugs, as it is also confirmed by the sharp increase in Nbs intended for cancer diagnosis and treatment^4^. In the last five years, the development of Nbs was strongly enhanced by the Sars-CoV-2 global pandemic. A wide number of Nbs were indeed produced as both a diagnostic and a therapeutic tool to face the global threat. Different targets were identified, the main one being the Receptor Binding Domain (RBD) of the Spike glycoprotein. During the infection onset, Sars-CoV-2 binds to the angiotensin converting enzyme 2 (ACE2) using the Receptor Binding Site (RBS), a subdomain of RBD^5^. Most Nbs developed to neutralize Sars-CoV-2 compete, partially or completely, with ACE2 for the binding to RBS.

The development of both Nbs and antibodies (Abs) is however time consuming and expensive. *In silico* methods^6^ have gained an increasingly relevant role in this process. Machine Learning (ML) and Deep Learning (DL) methods have been developed for the generation of sequences and for 3D structure prediction of Immunoglobulins (Igs), which entails both Abs and Nbs. However, Igs are more challenging to handle compared to other proteins, since CDRs, and CDR3 in particular, do not have an evolutionary conserved sequence which can guide the definition of rules and patterns for generative or predictive tools. This is because their sequences are generated through the *V(D)J* recombination occurring during the development of B and T cells, as well as through somatic mutations occurring during the so-called affinity maturation. This entirely random generative mechanism is responsible for the extremely diverse repertoire of CDR3 sequences^7^ which allows the immune system to recognise an incredibly diverse array of antigens. Identifying and characterising the complementarity-determining regions is therefore the key step in describing an immune repertoire, either in-vivo or during the synthesis of immuno-therapies.

The state-of-the-art methods to identify CDRs are based on similarities among immunoglobulin structures and sequences: some of the most widely adopted are Kabat^8^, Chothia^9^, AbM^10^ and IMGT^11^ numbering schemes. Each of these schemes identify CDRs from the sequence by searching for different recurrent amino acidic motifs. The fact that CDRs are generated randomly and only later selected based on their affinity with antigens represents however a massive difference compared to other sequence motifs such as transcription factor DNA binding sites or protein family domains. For these reasons, motif-based search methods applied to CDRs will inevitably show limitations, which are alleviated by successive updates of the definition rules as new molecules emerge.

Therefore, even though these schemes are largely reliable and widely used, inaccuracies, particularly in nanobodies, could potentially lead to confusion and discrepancies in the definition of CDRs^12^, and particularly CDR3. Structure prediction methods for Igs pose an even tougher challenge, given the peculiar malleability and conformational diversity of CDRs. Although tools such as AlphaFold3 (AF3) have enormously enhanced our capability to predict protein structures, they still have difficulties in predicting CDR3 conformations. Even though CDR3 loop prediction improves when the antigen is given, AF3 still has a 60 % failure rate in Abs and Nbs docking with their target when using a single seed sampling^13^. In summary, the intrinsic diversity of CDR3 sequences and conformations hampers the structure prediction and the docking precision.

Here, we asked whether we can harness the observed diversity/randomness in sequence generation and the apparent tendency for flexibility/disorder in immunoglobulin recognition regions to define a general, unbiased framework for CDR(3) identification. By employing a multi-disciplinary approach, we set out to address this key challenge using Nbs elicited against SARS-CoV-2 RBD as test cases. To proceed along this path, we make use of the general concept of fuzziness and fuzzy regions in the context of biomolecular interactions. From the sequence point of view, fuzziness in binding regions could aptly indicate a tendency for efficient exploration of the sequence space, free of constraints related e.g. to fold stabilization. From the structural perspective, fuzziness can favor adaptability to target partners via conformational selection mechanisms. This aspect has been associated with the function of intrinsically disordered regions (IDRs) involved in protein-protein interactions^14 15 16^.

We overcome motif-based or structure-prediction based methods for characterising the repertoire, which do not address the intrinsic biological characteristics of CDRs, and provide a straightforward method, based on deep-learning, to predict CDRs solely based on the Nb sequences. In parallel, we investigate the possibility to unbiasedly isolate a subset of surface residues from Nbs structures and recognize them as potential CDRs by analyzing their intraprotein energetic properties in the isolated, unbound Nb. Via the structure-based energy decomposition method, we indeed show that CDR3s displays a peculiar intramolecular interaction profile consistent with high intrinsic flexibility and consequent adaptability.

Taken together, our results define hypervariable loops as intrinsically disordered regions, demonstrate that conformational fuzziness underlies immunoglobulin polyreactivity, and show that independent sequence- and structure-based approaches converge on the same principle: explainable-AI methods reveal that the features learned by neural networks mirror those identified through interaction energy analyses, namely the intrinsically fuzzy nature of CDRs.

## Materials and Methods

### Development of a deep learning architecture

The model architecture has been developed and trained using a Docker container environment for reproducibility, with Python 3.11.0rc1 on linux and Tensorflow version 2.15.1, Keras version 2.15.0 running on hardware with NVIDIA-SMI 550.90.07, driver version 550.90.07 and CUDA version 12.4.

### Encoding and Decoding

The prediction of CDR sequences from a nanobody sequence has been designed as a classification task, i.e. predicting whether each residue in the sequence belonged to CDR1, CDR2, CDR3 or the rest of the protein (which we called “body”). Based on the annotated CDRs therefore, a script has been designed to search for each of the CDR sequences within the nanobody and convert each residue on its corresponding class, to be used as a label for the training of a supervised classification neural network. Body has been assigned class 0, CDR1 class 1, CDR2 class 2, CDR3 class 3. Additionally, to create a consistent dataset with equal-length inputs, each nanobody sequence has been padded to 150-residues length and padded null residues have been assigned class 4 (with a weight 0, for the training of the network). This scheme allows us to encode and decode any nanobody sequence and use the model predictions (see below) to extract the specific residues (sequence) of any CDR of interest in the nanobody sequence. As far as the input sequences are concerned, we simply assigned a sequential integer token to each amino acid residue, thus building a simple vocabulary of fixed size corresponding to all possible residues, plus a null-value residue for the padded regions.

### Architecture

Based on the encoding of choice, and the need to capture the context of the residues flanking each CDR on both sides, we decided to employ a sequential architecture tailored for sequence labelling tasks. The first layer of the network is an embedding layer (with dimension = 64), which serves the purpose of learning dense vector representations of the tokens from the above-described vocabulary, with a maximum input sequence length of 150 tokens. Following the embedding layer, we added two stacked bidirectional long short-term memory (BiLSTM) layers, with 32 and 64 units respectively, both configured to return full sequences in order to preserve the temporal context across the two layers. According to our choice regarding the encoding of the labels, the output layer consists of a time-distributed dense layer, with a softmax activation function, yielding class probabilities at each timestep across five categorical outputs (our categories), i.e. including the padding class. The model was trained using the Adam optimiser, and we imposed a learning rate of 1*10-4 and a gradient clipping (value 1.0), meant to mitigate potentially exploding gradients, when handling a large dataset encompassing millions of sequences. We used sparse categorical crossentropy as the loss function, and accuracy was computed comparing each residue on each nanobody sequence, with a weighting designed to ignore padding tokens (*Supplementary Fig. 1*). The total number of parameters in the model is 92.869, all of them trainable.

### Training and Validation data

The Integrated Nanobody Database for Immunoinformatics (INDI)^17^ is a comprehensive repository aggregating camelid single-domain antibody (VHH) sequences from five public sources: the Protein Data Bank (PDB)^18^ for structural chains, patent literature, NCBI GenBank records, high-throughput next generation sequencing (NGS) datasets and peer-reviewed publications. Each INDI entry consists of a complete VHH sequence linked to metadata from its original database. Raw sequences were filtered to obtain full-length VHH domains containing all three canonical CDR loops and comprising only the 20 standard amino acids. All sequences are annotated using an IMGT-based numbering scheme^11^. INDI comprises more than 11 million nanobody sequences. It provides the data in two linked flat-files (FASTA for sequences and tab-delimited metadata tables). This curated INDI dataset was used as the training corpus for our deep-learning CDR prediction model.

We split the data into 80% of the sequences used for training, and 20% of the sequences used for validation. In order to process such a large dataset efficiently, data have been chunked in tensors of 100,000 sequences each (i.e. shape [100,000, 150]), each saved as a separate file on disk. The training chunks were 90 containing a total of 8,982,880 nanobody sequences, while the validation chunks were 23, containing a total of 2,245,720 sequences. The total number of sequences used to build the model was therefore 11,228,600. We have trained the model for 4 epochs, each loading the data by chunk and limiting the batch size to 1024 to control the memory usage.

### Testing dataset

In this work, we selected a dataset of 3D structures of 121 nanobodies (Nbs) known to bind the Receptor Binding Domain (RBD) of the SARS-CoV-2 Spike glycoprotein. The Nbs were selected by downloading them from the Protein Data Bank^18^, and crossing information with that available in CoV-AbDab^19^, a Sars-CoV-2 dedicated database, and in SabDab-nano^20^, a nanobody centred repository. We extracted Nbs with corresponding 3D structures solved by X-Ray Diffraction Crystallography or Electron Microscopy, to generate a list of 121 non-redundant and monovalent biomolecules. Their respective PDB codes are reported in *Supplementary Table 1*.

### Computing infrastructure

The training of the model has been conducted using the Google Cloud Computing Platform. We employ Terraform^21^ to create reproducible infrastructure as-a-code (IaaC), and standardise our operations. To train the deep learning architecture on the INDI data, we configured a “a2-highgpu-1g” virtual machine, equipped with 12 vCPUs and 85GB RAM and a local SSD along with an NVIDIA Tesla A100 GPU with 40GB memory.

### Nanobody Characterisation

#### CDRs identification on Nbs sequences using Chothia

The *Multiple Sequence Viewer/Editor* tool from the *Maestro Schrödinger suite* version 2024-3 (*<u>Schrödinger Release 2024-3: Maestro, Schrödinger, LLC, New York, NY, 2024</u>*.) was employed to identify CDRs on Nbs by using the Chothia annotation scheme^9^. CDRs identified in this way are considered as the ground truth reference.

### Energy Calculation

#### Structure Preparation

The 121 selected Nbs structures downloaded from the PDB were first visually inspected via the Pymol molecular modeling package (*<u>The PyMOL Molecular Graphics System, Version 3.0 Schrödinger, LLC</u>*). Selected Nbs were extracted from the complex with RBD and crystallographic waters were removed.

The structures were then prepared for the energy calculation. First, residues missing from the original PDB structures were added. The *Protein Linker Design* tool from the *Maestro Schrödinger suite* version 2024-3 (*<u>Schrödinger Release 2024-3: Maestro, Schrödinger, LLC, New York, NY, 2024</u>*.) was used to reconstruct unsolved loops in five Nbs (7C8W_MR17, residues W111 and G112; 7CAN_MR17-K99Y, residues Y108, D109, Y110, W111 and G112; 7KN5_VHHU, residues G10 and G66; 7N9E_Nb34, residues D101, P102, Y103 and G104; 8BEV_W25, residues S55 and H56). The newly added residues form a linker between two attachment points. The *Interdomain link library* method was selected to predict linker conformations through a database of interdomain linkers from the PDB. The strain energy of the new loop was calculated in implicit solvent using *Prime*^22^ with the OPLS4 forcefield^23^, and it represents the difference between the linker in the minimized conformation adopted in the protein and in its free minimized conformation. The structure of the loop within the protein which maximises the difference to the free conformation, representing the minimum energy of the loop conformation in the protein, is selected as the representative conformation.

Next, each of the 121 Nbs underwent the *Protein Preparation Workflow*^24^ tool of the *Maestro Schrödinger suite* version 2024-3 (*<u>Schrödinger Release 2024-3: Maestro, Schrödinger, LLC, New York, NY, 2024</u>*.). Proteins were correctly protonated at pH of 7.4 and H-bonds assignments were optimized using PROPKA^25^, side chains missing from the original PDB were added and disulfide bonds between Cys residues with their respective terminal thiol groups within 3.2 Å were reconstructed. In the last step, Nbs structures were minimized using OPLS4 forcefield^23^ with the implicit solvent model implemented in Maestro.

#### Energy Decomposition Method for the Prediction of Potential Interacting Regions (CDRs) on Nanobodies

The structures of the Nbs prepared as described above were subsequently converted into an Amber-compliant PDB file using the *pdb4amber* utility included in the *Amber24 suite*^26^.

Next, each Nb structure was used as input for the structure- and interaction energy-based prediction of potential interacting regions using the Matrix of Low Coupling Energies (MLCE) method^27 28 29^.

The scripts and instructions to run this analysis are available on GitHub (https://github.com/colombolab/MLCE). Proteins are prepared and analyzed using the AmberTools suite^26^, following the steps described below.

First, each Nb PDB is converted to the AMBER format by using the *tleap* utility, obtaining a topology file and a configuration file (respectively in.*prmtop* and.*inpcrd* format). Next, the system is minimized in implicit solvent through 200 steps of steepest descent minimization using the *sander* executable.

In the third step, the non-bonded part of the potential, including van der Waals, electrostatic interactions and solvent effects, is computed via an MM/GBSA calculation performed with AmberTools python script *MMpbsa*.*pl*. For a Nb of *N* residues, the output is a *NxN* symmetric interaction matrix *M*_*ij*_, in which only van der Waals and electrostatic terms are summed:

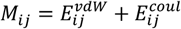

The original Energy Matrix can be diagonalized and reconstructed using the resulting eigenvalues and eigenvectors:

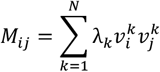

where *λ*_*k*_ is the *k*th eigenvalue, and 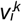 is the *i*th component of the corresponding eigenvector. The eigenvectors are then labeled from the most negative to the most positive. We previously showed^27 28 29^ that it can be assumed that the first (most negative) eigenvalue, labeled *λ*_*1*_, and the associated eigenvector, namely ***v***_***1***_, contain most of the information on the stabilizing interactions of the system. An approximated interaction matrix 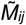 can thus be defined as:

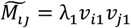

As the 3D structure of the protein under exam is known, for each Nb a contact matrix ***C***_*ij*_ can be built by considering two amino acids in contact if the distance between two of their heavy atoms is smaller than 6Å threshold. The Hadamard product of the two matrices gives us the matrix of the local coupling energies, i.e. information on the pairs of residues in contact in the 3D structure and the energetic intensity of this coupling.

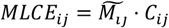

Potential interaction substructures are identified as sets of close-by residues that show weakest coupling interactions with the rest of the protein. Regions of weak coupling, also called patches, are thus identified by setting a percentage threshold. In this work, a 10% threshold was used, meaning that patches are composed of residues involved in the 10% of interactions that are less favourable energetically.

In this framework, regions of weak coupling with the rest of the protein also coincide with substructures that are generally prone to undergo substantial conformational changes, tolerate mutations without impacting on the overall 3D organization of the cognate biomolecule, and adapt to a binding partner with minimal energetic expense. This approach was validated in multiple instances against experimental data^28 30^. The aforementioned properties can also be considered hallmarks of fuzziness for putatively interacting regions.

In the case of Nbs, it can be aptly hypothesized that the regions that emerge as most prone to interaction may in fact coincide with Complementarity Determining Regions (CDRs). As it is shown in *Supplementary Table 1*, patches identified on Nbs include residues of CDRs in most cases.

### Explainability

On the trained model used for inference on the RCSB dataset, different methods have been used to explain the predictions, described in the following paragraphs.

#### Saliency analysis

Gradient-based saliency maps were computed to assess the relative importance of each input position for model predictions across sequence timesteps. For each input sequence in inference mode, we identified the predicted class at each output timestep and then computed the gradient of the corresponding class probability with respect to the embedding output. This has been done using TensorFlow’s automatic differentiation (*tf*.*GradientTape*). The model was functionally decomposed into its embedding layer and downstream recurrent layers, enabling position-wise gradient analysis. At each timestep *t*, the absolute gradient values across embedding dimensions were averaged to yield a saliency score for each input position *p*, resulting in a full saliency matrix indexed by both timestep and position. These values were initially recorded as individual records for every (*t, p*) pair. To facilitate interpretability, we then aggregated the saliency values by averaging across *t*, producing a summary measure that quantifies the average importance of each residue *p* in predicting class *c* at any position in the sequence. This final representation allows position-wise comparison of contributions to model decisions independent of output location. We used these values to produce saliency plots for each of the predicted classes along each nanobody sequence.

#### Hidden states analysis

Hidden state vectors from the final bidirectional LSTM were extracted for all non-padding residues across all input sequences of the RCSB dataset described above. The trained model was decomposed to expose the intermediate activations: this was done by assembling a sub-model from the original input and through the embedding and following recurrent layers, but terminating prior to the classification output. For each sequence we then computed the full sequence of hidden states and collected those corresponding to non-padding positions. Hidden states were stored alongside their metadata including the input position, predicted class at that position, token identity and sequence identifier in order to facilitate downstream analysis.

Principal component analysis was then applied to the matrix of hidden state vectors, after normalising all numeric features. PCA was performed using the *tidymodels* package, and projecting the data on 2 principal components to facilitate visual representation. We then joined the resulting dataset with residue-level metadata, including the measure of energy we were interested in as well as physicochemical attributes, such as thermodynamic β-sheet and coil propensities^31^ for comparison.

### Data availability

The data used to develop the deep learning model are available on INDI^17^, and the data used to carry out the analysis are available on PDB, with the identifiers mentioned in Supplementary Table 1. The code is openly accessible and documented on GitHub at https://lescailab.github.io/nanocdr-x/ and available as a conda package at *lescailab::nanocdr-x*.

## Results

### Deep learning model performance

The architecture (*Supplementary Fig. 1*) trained on INDI^17^ achieved an accuracy higher than 0.995 already at 2 epochs, with minimal but steady increase until 4 epochs. No sign of overfitting can be reported, since the validation accuracy maintained values above the training accuracy until epoch 4 where the training was stopped (*Supplementary Fig. 2*).

The final validation accuracy reached 99.9 % on the INDI dataset. The model thus trained was therefore saved for distribution along with the encoding and the decoding scripts, as well as interpretability tools, which we discuss later.

The software is distributed with a conda package easily accessible at *lescailab::nanocdr-x* and reference documentation has been made available at https://lescailab.github.io/nanocdr-x/ where the source code can also be accessed.

A perfomance test at first installation showed that the analysis of 121 nanobodies (indipendent dataset, see below) required 43 seconds of wall-clock time (real 0m43.061s) with a user CPU time of 5.696s.

### Independent dataset validation

We selected an independent dataset, with experimentally resolved structure, with the goal of: (1) assessing the model accuracy beyond the training and validation data and (2) comparing the predictions with consolidated methods in the literature such as Chothia^9^. In order to do so we used the CDRs predictions resulting from the Chothia (annotated with MAESTRO tool) scheme as the ground truth, and encoded each residue in a class number in the same way as our model classification detailed in *Materials and Methods*. To each position in the sequence we assigned class labels as follows: 0 for body (defined as any nanobody sequence position not classified as CDR), 1 for CDR1, 2 for CDR2 and 3 for CDR3. Padding positions were excluded from the comparison.

We then compared the class annotated with Chothia with the class predicted by nanocdr-x on each residue of each nanobody and generated the summary results as shown in Table 1 (confusion matrix). The results show strong concordance between the predicted and reference loop regions. The majority of predictions fell along the diagonal of the confusion matrix, indicating a high level of class agreement. Most notably, class 1 (CDR1) showed 835 correctly predicted residues out of 835, class 2 (CDR2) was predicted for 695 residues out of 695 and class 3 was predicted for 1697 positions out of 1,699 classified by Chothia as CDR3. As far as no-CDRs sequences (body) nanocdr-x predicted 11097 correctly classified residues out of 11,758. Overall this analysis has revealed our model to yield an overall F1-accuracy score of 95.6%.

**TABLE 1.** Confusion matrix comparing Chothia and nanocdr-x loops encodings. Each cell reports the number of residues assigned to a given class by Chothia (rows) and by nanocdr-x (columns). Values along the diagonal indicate concordant classifications, whereas off-diagonal values correspond to discrepancies.

| Confusion Matrix |  |  |  |  |  |
| --- | --- | --- | --- | --- | --- |
| True Label<br>(Chothia) | Class 0 | 11096 | 140 | 249 | 273 |
|  | Class 1 | 0 | 835 | 0 | 0 |
|  | Class 2 | 0 | 0 | 695 | 0 |
|  | Class 3 | 2 | 0 | 0 | 1697 |
|  |  | Class 0 | Class 1 | Class 2 | Class 3 |
|  |  | Predicted Label<br>(nanocdr-x) |  |  |  |

### Benchmarking

To evaluate the performance of our tool (nanocdr-x), we conducted a comparative analysis with the main tools described in the literature which mentioned CDR loop prediction in either antibodies or nanobodies. The comparison focused on approaches using deep-learning, with specific attention to those explicitly designed to handle nanobody sequences or to predict the CDR loops. The final selection of tools was guided by a systematic review of recent literature^32^ and includes: simpleDH3^33^, AntiBERTa^34^, AntiBERTy^34^, ProtBERT^35^, nanoBERT^36^, ABlooper^37^. However, none of these tools provided an immediate way to extract the CDR loops from the primary sequence alone.

Several approaches relied on three-dimensional structures. Most of the tools we analysed, rather than predicting which residues of a known nanobody sequence belonged to a CDR loop, carried out a sequence prediction of unknown (masked) residues within a sequence. In our benchmark, we therefore realised that most of the tools we tested did not carry out a straightforward sequence classification, and that CDRs identification could not be carried out as we intended.

Most of these approaches, however, were developed for different purposes: some rely on three-dimensional structural information, while others are primarily aimed at predicting masked residues within sequences rather than performing a direct classification of residues into CDR and non-CDR regions. As a consequence, a direct comparison at the level of sequence-based CDR annotation was not always feasible.

In contrast, nanocdr-x was specifically designed to perform an end-to-end classification of nanobody sequences, assigning each residue to one of the four classes (body, CDR1, CDR2, CDR3) directly from the primary sequence.

### Characterising the binding loops

#### Structure-Energy based identification of CDRs (paratope prediction)

In parallel, we perfomed the prediction of CDRs on a database of 121 non-redundant Nbs known to bind the Receptor Binding Domain of Sars-CoV-2. As we reported in *Materials and Methods*, we chose Nbs on the basis of the availability of their experimental three-dimensional structure solved by either X-Ray Crystallography or Electron Microscopy and deposited in the Protein Data Bank. After structure preparation (see *Materials and Methods*), we predicted paratope regions on each Nb structure in isolation using our in-house method MLCE (Matrix of Low Coupling Energy^27 28 29^). Our working hypothesis is that CDRs need to have a peculiar energetic signature, allowing them to adapt to the binding partner via significant conformational changes and sampling of diverse structural populations, from which the binding states can be selected. This type of behavior identifies CDRs as fuzzy loops. Therefore, it is reasonable to hypothesize that their residues are weakly coupled with the rest of the protein from the energetic point of view. The formation of a localized network of weak interactions is in fact a distinctive trait of fuzzy regions. As a consequence, to identify them, we used an energy-pair decomposition approach, namely MLCE.

MLCE analyzes the interaction energies between all amino acid pairs within a protein with a solved 3D structure. First, an MM/GBSA calculation is performed to compute the nonbonded part of the potential, *i*.*e*. electrostatic and van der Waals interactions and solvent effects, yielding an *N* × *N* symmetric interaction matrix *M*_*ij*_ for a protein of *N* residues. The matrix is then diagonalized and reconstructed through eigenvalue decomposition to identify regions of strong and weak energetic coupling. Since it can be assumed that the first (most negative) eigenvalue and the associated eigenvector contain most of the information on the stabilizing interactions of the protein (see *Materials and Methods*), we then analyzed the contribution of each residue to the most negative eigenvector. As an example, values calculated for Nb7A29 are reported on the y axis (energy) in the plot in figure 2A, while residue numbers are listed on the x axis. Highest energy values correspond to a high contribution to the most negative eigenvector and are characteristic of stabilizing regions of Nbs (*i*.*e*. the folding core or stability core), while lowest values are distinctive of weakly coupled regions, normally associated with PPI surfaces. We consistently identified regions with low values by setting a threshold for each Nb calculated as 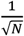. This limit represents an ideal situation in which each of the *N* residues contributes equally to the first eigenvector (flat eigenvector hypothesis). Consequently, residues with higher energy values compared to the threshold are important for the stabilization of the protein fold and their energetic contribution is represented as blue bars in figure 2A, while residues with lower energy values belong to regions with more conformational variability, and thus possibly important for the interaction with partners. CDR1/2/3 have the lowest values in the plot and their corresponding bars are depicted in red. The contribution of each residue to the first eigenvector can be projected on Nb7A29 structure, as shown in figure 2A-E: the color spans from blue (stabilizing regions) to red (weakly coupled regions). This trend, consistently observed over the 121 Nbs we analyzed, indicates that CDR regions always correspond to residues weakly coupled to the rest of the protein. In the last step of the MLCE procedure, the interaction matrix *M*_*ij*_ is filtered by the contact matrix *C*_*ij*_ (*see Materials and Methods*) in which potential interacting patches are rigorously identified as sets of close-by residues weakly coupled with the rest of the protein. These characteristics are always typical of CDRs, as they are important interacting regions, and can also be considered as hallmarks of fuzziness. In this context, we envision that calculations like the ones presented here could aptly be used to predict CDRs from simple structural information.

Importantly, patches identified on Nbs often include residues belonging to CDRs according to Nanocdr-x predictions (*see previous paragraphs*). In *Supplementary Table 1*, the results of our MLCE-guided paratope prediction on Nbs show that in 112 out of 121 Nbs the predicted patches include one or more CDR3 residues (90,91 % success). As an example, in figure 1A interacting regions predicted using MLCE are highlighted on the structure of 7KKK_Nb6: as it can be observed, CDR loops are correctly included in the patches. In figure 1B, a boxplot reports the distribution of interaction energy values across body residues and the three CDRs. A one-way ANOVA confirmed that the differences among residue classes are highly significant (F(3, 14,983) = 682.6, p < 2.2×10^−16^), supporting the hypothesis that CDR loops correspond to weakly coupled, energetically uncoupled regions when compared to the structural core of nanobodies.

**Figure 1.**
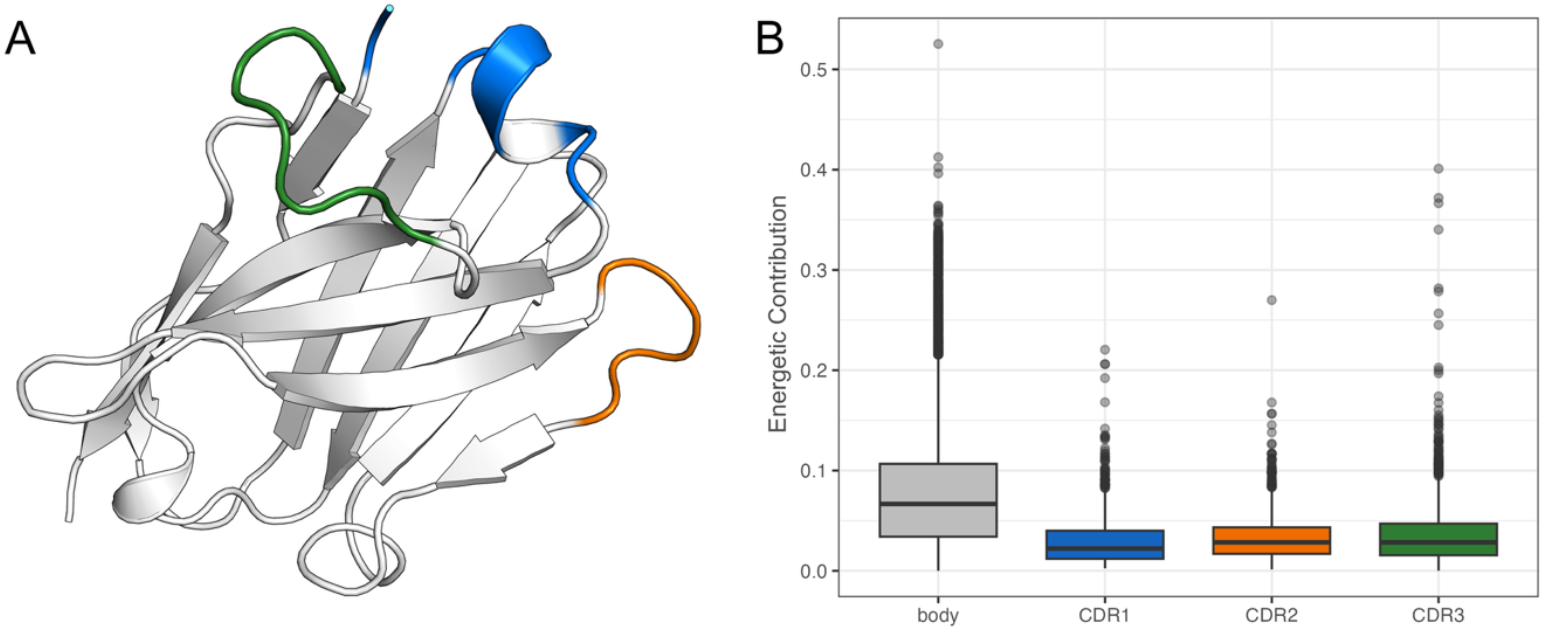
Structure- and energy-based identification of CDRs. (A) Predicted interaction regions using MLCE on the structure of nanobody 7KKK_Nb6 (PDB ID: 7KKK). The protein fold is shown in cartoon representation (grey), with predicted interacting patches in orange, blue, and green. These respectively correspond to CDR1, CDR2, and CDR3, showing that MLCE correctly identified CDRs as weakly coupled, flexible regions.(B) Distribution of interaction energy values for residues classified as body or CDRs. Boxplots show significantly lower energy values in CDR1–3 compared to body residues (ANOVA, F(3, 14,983) = 682.6, p < 2.2×10^−16^), indicating that CDRs consistently display a characteristic energetic signature of weak coupling.

#### Explainability

The excellent performance of our model prediction addresses one of the main challenges we sought to resolve, i.e. making CDRs predictions accessible and quick. However, this accuracy solely based on sequence context opened an additional opportunity to better understand the molecular and biochemical characteristics of those regions. The structural characterisation of CDRs obtained using the energy decomposition method presented in the previous paragraph revealed the fuzzy nature of these regions. We therefore analysed the performances of our model in terms of saliency and hidden states to capture the internal representation of Nbs and the relationship with structure-based energetic calculations.

#### Saliency

Firstly, we focused on the relationship between saliency and energy measures. Saliency is computed (see *Materials and Methods*) on the model gradient, to assess the average importance of each residue in allowing the model to predict each class. Figure 2A shows the values of the energy measured along nanobody SB-23 in structure 7A29; figure 2B plots the importance of different residues in the prediction of CDR 1 positions; figure 2C plots the importance of each residue in achieving the prediction of the CDR2, and figure 2D plots the average importance of residues in allowing the model to identify CDR3. The three plots combined show that the most important positions for the neural network to identify the binding loops correspond to areas of low energy, and only in part to framework positions. A central element of the saliency analysis is that the highest importance in predicting each loop seems to reside in the loop itself, rather than in surrounding regions. This is significantly different from the established methods in the literature, which cannot identify randomly generated sequences such as the binding loops, and therefore focus on identifying loose patterns in the surrounding regions. Figure 2E provides a structure-based context to these observations. The plot shows the importance of the loops themselves in predicting the binding regions, together with flanking sequences which, according to their energy measures, might contribute to the stability of the molecule.

**Figure 2.**
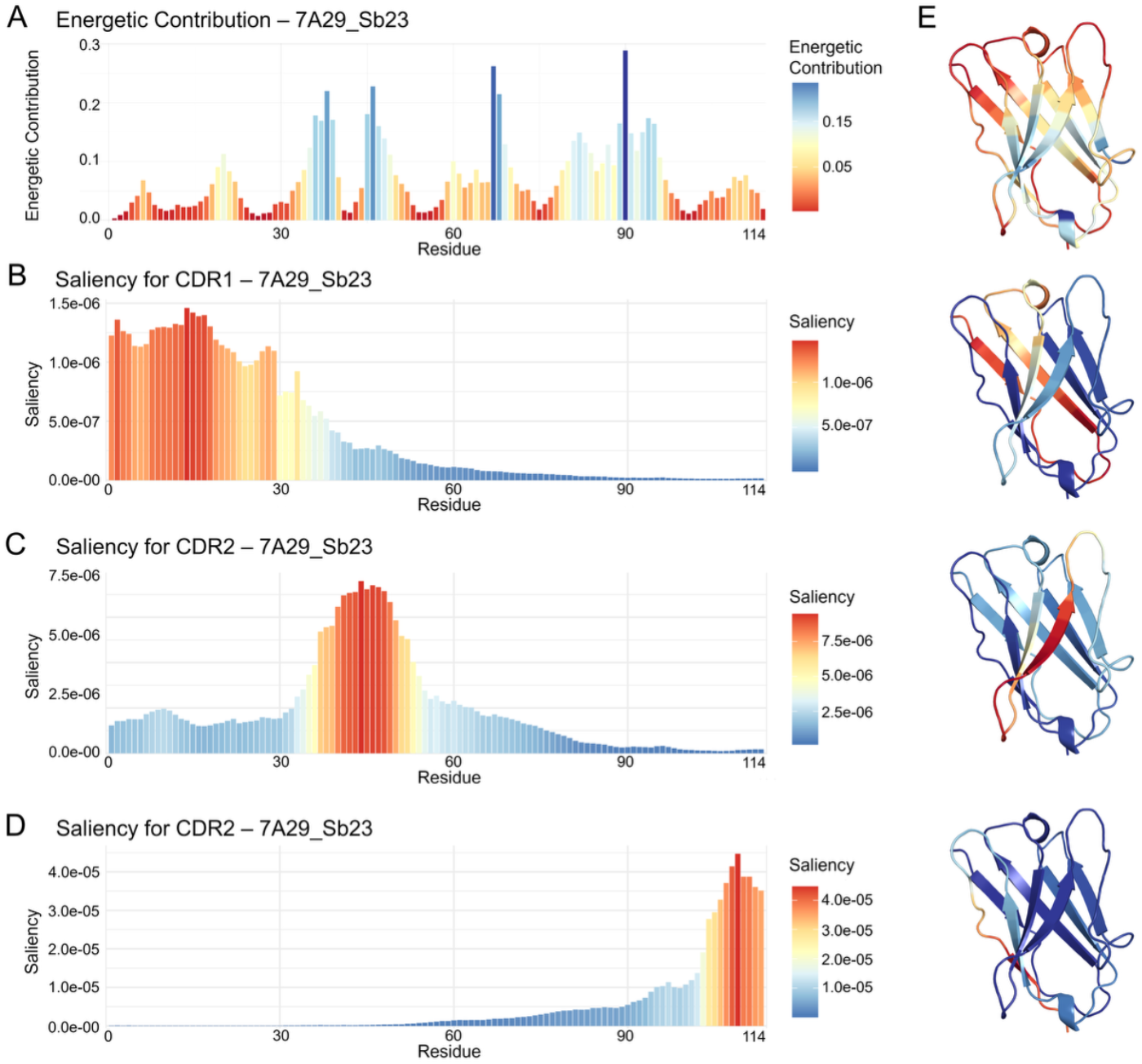
Saliency mapping of nanobody 7A29_Sb-23 (PDB: 7A29) in relation to structural energy. Residue-level saliency(2B,2C,2D) scores, derived from model gradients, indicate the importance of each residue for CDR classification. Regions with high saliency overlap with low-energy(2A) residues of the structure and are concentrated within the CDR loops. This alignment highlights the model’s ability to focus on structurally flexible yet functionally relevant areas. Compared to state-of-the-art methods, the approach achieves improved localisation of CDRs, with predictions supported by structural stability mapping. The right panel (2E) shows the projection of values from the respective plots (2A, 2B, 2C, 2D) on the 3D structure of 7A29_Sb-23

#### Hidden states analysis

To further understand if the molecule flexibility was indeed what the architecture has captured, we decided to analyse the internal representation of the nanobody, learned by the model. This is done by extracting the numerical representations (hidden states) of each residue built by the last layer of the network, before the classification output. Given the dimensionality of the last LSTM layer, each aminoacid is represented by a vector of 128 values. Therefore, dimensionality reduction was used to facilitate the interpretability and visualisation of such representations, and assess any relationship with the energy measure as well as with other residue characteristics annotated from the literature. We performed a principal component analysis (PCA) of the hidden states thus extracted, evaluating the distribution of each nanobody position along the first two principal components. The PCA plot shows a clean separation of the different CDRs on the PCs plane, thus confirming the ability of the model to separate (i.e. predict) the different classes.

We then added the energy measure as a third dimension, to better assess the relationship not only with the prediction (saliency) but also with the model representation of the nanobody (hidden states). Figure 3A shows that the denser areas of the residues predicted as the three CDRs correspond to lower energy regions. Alternatively, adding the thermodynamic beta-sheet propensity^31^ as a third dimension (Figure 3B) shows a randomly scattered distribution of the very same residue representations. The comparison of the two annotations shows that the classical thermodynamic structure propensity measures do not cluster with the CDRs representations, while most clearly the fuzziness of the molecule does.

**Figure 3.**
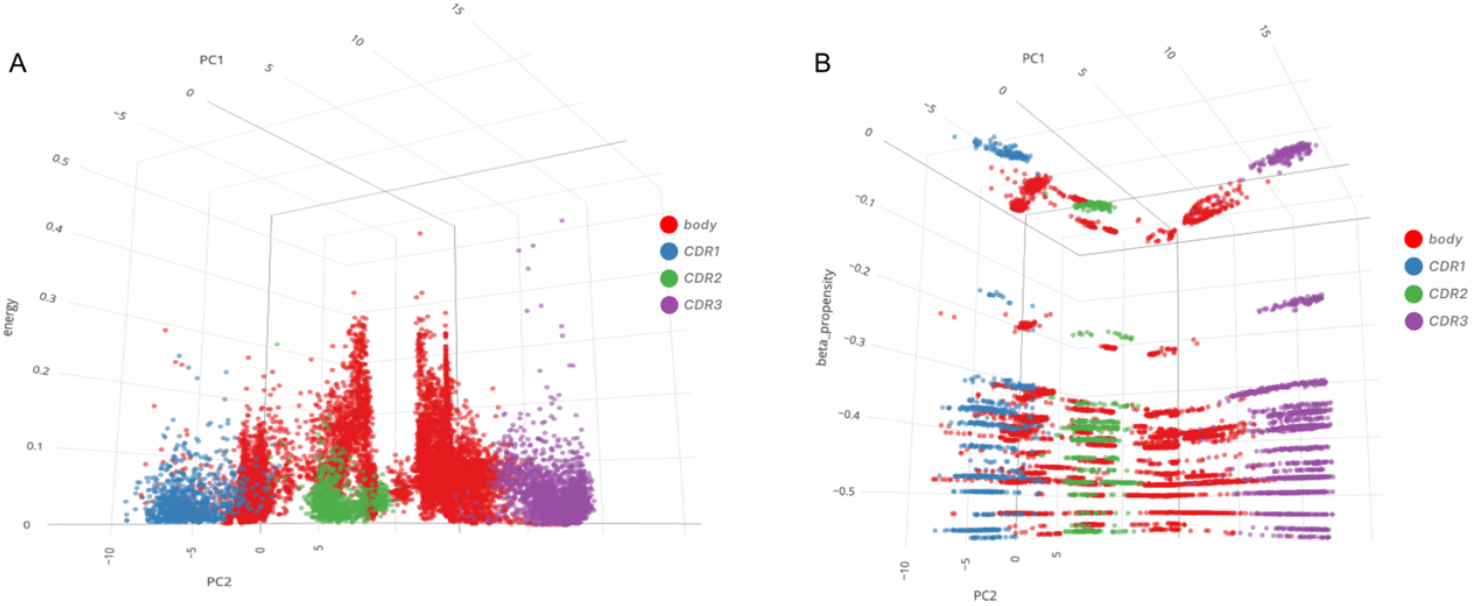
Projection of hidden state representations onto structural properties. Hidden states from the neural network capture how the model represents nanobody sequences. When projected onto β-sheet propensity values (B), the embeddings appear evenly scattered and do not separate CDRs from framework regions. In contrast, projection onto structural energy decomposition (A) results in distinct clustering that aligns with CDR classification. This suggests that the features the model learnt align with the intrinsic “fuzziness” of nanobody regions, as reflected by their energy scores, thereby supporting accurate loop identification.

## Discussion

Nanobodies (Nbs) have emerged as an interesting class of antigen-binding biomolecules, with applications that span from next-generation therapeutics to precision diagnostics. Nbs are relatively small single chain molecules, which makes them versatile tools easier to produce and handle compared to larger multi-chain antibodies.

Understanding the determinants of their evolution and of selective antigen binding is crucial to rationally engineer novel nanobodies. In this context, investigations have long concentrated on clarifying antigen-binding mechanisms through structure-based methods, which rely on molecular docking, homology modelling, and most recently on the use of AI-driven approaches, such as AlphaFold2 and AlphaFold3^13^.

A first critical step toward uncovering the key determinants of Nbs mechanisms, and possibly developing rules for their design, resides in the unbiased identification of Complementarity-Determining Regions (CDRs), i.e. those loops responsible for the interaction with the antigen. The key question is whether a unifying framework can be developed integrating sequence and structural properties to capture the emergence of the unique functional features that characterize CDRs residues.

Yet, characterising CDRs, especially CDR3, remains a key challenge: their sequences arise from an inherently random mechanism resulting from the *V(D)J* recombination, as well as from further somatic mutations arising during the affinity maturation process. The structure of these loops tends to be flexible and disordered, and predicting their localisation may turn out to be challenging even for state-of-the-art methods looking at the 3D context such as AF3.

In this work, we set out to define a general and unbiased framework for CDR identification, simultaneously addressing both sequence and structure. The aim is to define a general signature that connects such properties while explaining the intrinsic specificity of CDR regions in antigen recognition.

From a sequence perspective, we worked on the hypothesis that using predefined schemes for CDRs identification was in (strong) contrast with the intrinsic variability of CDRs sequences, and in particular of CDR3. We developed **nanocdr-x**, a deep learning architecture trained on more than 11 million nanobody sequences. For the first time, a model could robustly predict CDRs sequences and their localisation, with high accuracy. This approach fills a methodological gap: nanocdr-x is the only available tool that provides an end-to-end, open source, and easily accessible solution for large-scale automatic CDRs identification directly and solely from sequence data.

Importantly, this result focused our attention on the molecular properties of nanobodies and what the model had learnt. nanocdr-x does not need any additional information, such as structure or residue annotations. The saliency analysis, i.e. measuring the importance of each residue for the model to achieve its prediction, placed the highest saliency values within the CDRs themselves. This could only be interpreted with the existence of specific properties which are intrinsic to the CDR residues, and which the neural network could uniquely recognise to achieve the successful prediction of the CDR sequence and localisation in the protein.

We next asked whether sequence features may propagate into structural properties characteristic of CDRs. CDRs are known to sample diverse conformational ensembles, from which a preferred state is selected upon antigen binding. Such a conformational selection mechanism parallels the behavior of intrinsically disordered regions, enabling efficient recognition and adaptability to the target. We reasoned that this flexibility arises when residues within CDRs are only weakly coupled to the Nb global fold, allowing them to adopt multiple conformations and accommodate high mutational variability without compromising the protein’s structural integrity. Antigen engagement would then stabilize otherwise dynamic residues, favoring the formation of a functional complex.

To identify CDRs from a structural standpoint, we applied the Matrix of Low Coupling Energy (MLCE) method^30^, which analyzes intramolecular interaction energies across all residues of a protein with an available structure. MLCE assumes that interacting patches are defined by suboptimal intramolecular energetic pair-interactions, conferring flexibility and adaptability to binding partners. Using only the 3D structure of isolated nanobodies as input, MLCE consistently mapped CDRs as weakly coupled regions, without requiring prior knowledge of sequence motifs. These loops emerged as energetically decoupled from the protein core, supporting the view that low coupling underpins their conformational plasticity, capability to support high mutation rates, and functional adaptability.

Strikingly, two completely independent methodologies pointed to the same conclusion, prompting us to investigate how they could be interpreted in light of each other. The explainability analysis of the deep-learning model clearly showed that the intrinsic properties recognised by the neural network in CDRs directly corresponded to the low-energy values identified by MLCE. Unsurprisingly, when testing nanocdr-x explanations with conventional residue properties such as beta-sheet propensity, they were not able to account for the predictions achieved by the model.

In light of these findings, the concept of fuzziness emerged as the unifying hallmark characteristic of CDR regions in nanobodies. We speculate that this concept can be aptly extended to the more general realm of antibodies.

In this context, an emerging area of literature is increasingly focused on intrinsically disordered regions (IDRs) involved in protein-protein interactions^14^. These particular substructures are disordered in isolation and select optimal configurations upon binding to their partner. This feature has been shown to play a key role in recognition processes across different scales^38 16^. A relevant example in this context is represented by the formation of the chemokine heterodimer between CXCL12 and frHMGB1^15^. The realization that fuzzy regions are widespread in protein mechanisms also recently attracted attention for drug development^39^.

From our results, we propose a model where CDRs are inherently fuzzy: in this picture, fuzziness is a key feature in determining Nbs polyreactivity, since it can efficiently facilitate adaptation to different antigens. Since CDRs rigidify upon affinity maturation^40^ paratope fuzziness may be the decisive feature determining which Nbs (and potentially antibodies) progress to this stage, before rigidity is imposed to enhance the affinity for a specific epitope, via the acquisition of mutations that energetically stabilize the complex.

Fuzziness is not noise, it is function. Chaos here is adaptive. Disordered regions are all but random: their fuzziness is indeed the key property which confers the necessary adaptability to be selected as the most effective binding protein to an antigen target.

This work highlights the power of interdisciplinarity, i.e. combining computational genomics and computational chemistry, to reveal new biological principles. Beyond this conceptual advance, it paves the way for rational in-silico design strategies for antibodies and nanobodies.

## Acknowledgements

This work was supported by the grant of the Ministry of Health: “Piano Operativo Salute – Traiettoria 4, per la creazione di hub delle scienze della vita”, project “Immuno-Hub”

## Authors contributions

FL and GC conceived the work; DB and GT carried out the analyses; FL, GC, DB and GT wrote the manuscript; FL and GC revised the manuscript.

## Competing interests

The authors declare that the research was conducted in the absence of any commercial or financial relationships that could be construed as a potential competing interest.

## Notes

### Competing Interest Statement

The authors have declared no competing interest.

