## Supplementary Information for "Adaptive disorder as the hallmark of nanobodies antigen-binding loops"

**Supplementary Table 1 | Predicted interacting residues and CDR assignments for the 121 nanobody structures analyzed**

| PDBID_Name | CDR1_residues | CDR2_residues | CDR3_residues | S |
|---|---|---|---|---|
| 6YZ5_H11-D4 | 26GRTF29 | 54SGGS57 | 104SLLS107 – 111TWP113 | YES |
| 6ZCZ_H11-H4 | 25GRTF28 | 53SGGS56 | 99HYVSYLLSD107 | YES |
| 6ZXN_Ty1 | 25GF26 | 53PNSG56 | 101NL102 – 104SS105 | YES |
| 7A29_Sb23 | 26GFP28 | 53STGGW57 | 100VGY102 | YES |
| 7B27_NM1230 | G26 | 54AGDL57 | 102RVGTGER108 | YES |
| 7C8V_SR4 | 26GFP28 – 30YS31 | 54HGD56 | V100 | YES |
| 7C8W_MR17 | 26GF27 | -- | 104LA105 | YES |
| 7CAN_MR17-K99Y | 26GF27 | -- | 101DGQLAYHYDY110 | YES |
| 7D2Z_SR31 | 29GFP31 – 33WQ34 | 56SMGYK60 | 103VG104 | YES |
| 7F5G_DL4 | 27GSFFE31 | -- | 100EGQR103 | YES |
| 7F5H_DL28 | 26GSDFSS31 | 52SEGS55 | D98 – W100 | YES |
| 7FAT_Nb1A7 | 26GYT28 – S31 | S53 | 99GGPSLSYCTGG109 – 11GFL113 | YES |
| 7FAU_Nb1B11 | 25GYT27 – 29SV30 | 52ASGI55 | 97GLVRGSCTD105 – Y112 – G114 | YES |
| 7FBJ_17F6 | 29SCA31 | -- | S88 – 90SAGMCA95 | YES |
| 7FBK_20G6 | 29PCS31 | -- | 90TGDER94 | YES |
| 7FG3_K-874 | 26GS27 | 55GGT57 | 100GGDG103 | YES |
| 7JVB_Nb20 | 26GAGA29 | 51ASGG54 | 99IETA102 | YES |
| 7KGJ_Sb45 | G26 | 54AGQ56 | 102GHHYE106 | YES |
| 7KGK_Sb16 | 26GFP28 – 30AY31 | 54YGIK57 | -- | NO |
| 7KKK_Nb6 | 25GII27 – 29GR30 | 52RRGSI56 | 100ASPAPGD106 | YES |
| 7KLW_Sb68 | 29GSISSI34 | 56TVNGH60 | A103 – 105GY106 | YES |
| 7KM5_Nanosota 1 | 26GFT28 – 30KN31 | -- | 98GSKSGHEL105 | YES |
| 7KN5_VHHE | 26GVTLDY31 | -- | 101GTYYSGNYH109 | YES |
| 7KN5_VHHU | 25GF26 | 52SSGG55 | 99SGSYY103 | YES |

| | | | | |
|---|---|---|---|---|
| 7KN6_VHHV | 26GF27 | -- | 100SLGGWG105 | YES |
| 7KN7_VHHW | 25GFT27 | 52SSGD55 | 101SYYW104 | YES |
| 7LX5_WNb10 | G26 | -- | 104TWPE107 | YES |
| 7LX5_WNb2 | 25GFTLDY30 | 53SGGN56 | 100ATYYSGSYY108 | YES |
| 7MDW_Nb105 | -- | -- | 103WG104 | YES |
| 7MEJ_Nb36 | -- | -- | -- | NO |
| 7MFU_Sb14 | 26GFPVQA31 | 53STGT56 | 100VGSS103 | YES |
| 7MY2_Nb30 | 25GLT27 | -- | -- | NO |
| 7MY3_Nb12 | G26 – 29HN30 | -- | 100PYFGNSCV107 | YES |
| 7N9B_Nb21 | -- | 51ANGGN55 | -- | NO |
| 7N9C_Nb95 | 26GR27 | -- | 102VYY104 | YES |
| 7N9E_Nb34 | 25GF26 | -- | 99KDPYGSPWT107 | YES |
| 7N9T_Nb17 | 26GSI28 | -- | 101SAYAP105 | YES |
| 7NKT_NM1226 | 26GSLDY30 | 52SSGD55 | 98LQGSYYY104 | YES |
| 7NLL_Fu2 | 26GF27 – D31 | 54SDG56 | 100PSFSYTGSTYY110 | YES |
| 7OAO_C5 | 26GVTLGR31 | F54 | 100VTAA103 – S105 | YES |
| 7OAP_C1 | 26GFTNDF31 | 54SDNT57 | 101FAG103 | YES |
| 7OAP_H3 | 26GRT28 – 30ST31 | 54TGSS57 | 100TIV102 | YES |
| 7OAY_F2 | 26GRT28 – S31 | 53WSSTP57 | 101GESYY105 – R108 | YES |
| 7OLZ_Re5D06 | 26GITLD30 | -- | 100PLTKYGSSWY109 – P111 | YES |
| 7OLZ_Re9F06 | 26GRTF29 – N31 | 53WNSG56 | 99SDGYL103 | YES |
| 7P77_Sb-15 | 26GFPVKN31 | 53SGGV56 | 100VGR102 | YES |
| 7Q3Q_VHH-12 | 26GLT28 – S31 | 52RWKFGN57 | 101VG102 – 105IAV107 | YES |
| 7Q3R_VHH-F04 | 25GRA27 | 53SGGS56 | 99VDYSGTLTAA108 | YES |
| 7Q3R_VHH-G09 | 26GTGFT30 | -- | 100FDSSDYEVL108 | YES |
| 7R4I_Nb2.15 | 26GYASWAR32 | 54DFDG57 | 102GT103 | YES |
| 7R4Q_Nb1.29 | 26GYTINT31 | 54GSGN57 | 101YGASGYD107 | YES |
| 7R4R_Nb1.10 | 26GYTYSTC32 | 53ADG55 | 99VKDFT103 – T105 | YES |
| 7RBY_Nb112 | 26GLTLDY31 | 54SDG56 | 100PSTYYSGTYY109 | YES |
| 7TPR_7A3 | 26GYTSSS31 | -- | 102YNQWG106 | YES |
| 7TPR_8A2 | -- | -- | 101TYDKYAPCGGFAGTY115 | YES |
| 7VNB_n3113 | -- | -- | 102GSTG105 | YES |

| | | | | |
|---|---|---|---|---|
| 7VNE_n3113.1 | -- | 52PSGR55 | 100SGSTG104 | YES |
| 7VOA_aRBD5 | 26GF27 | 54SGGI57 | 101HTVVAGC107 – 112WTDF115 | YES |
| 7VQ0_P86 | 26GR27 | -- | 97DVNGGM102 | YES |
| 7W1S_Nb007 | 25IS26 – 28SSF30 | 50GIGG53 | -- | NO |
| 7WD1_R14 | 26GFTLDY31 | 54SDG56 | 100PATYYSGRYYYQ111 | YES |
| 7WD2_S43 | 26GFT28 – Y31 | 52SSN54 | 99PDYSGVYYYT108 | YES |
| 7WHI_Bn03nano 1 | 26DSS28 | 53PSG55 | 102GST104 | YES |
| 7WHI_Bn03nano 2 | D26 | 53GLGGA57 | 102FG103 | YES |
| 7X2J_Nb70 | 26GRT28 | 54DGGT57 | 100GNQYYSAT107 – S109 | YES |
| 7X2L_3-2A2-4 | 26GSISTL31 | 52TLDGS56 | 100GGF102 | YES |
| 7X2M_1-2C7 | 26GD27 | 53PSGSR57 | 101PSAHY105 | YES |
| 7X4I_aSA3 | 26GFTSDH31 | S53 – 55GNP57 | 99LWYGR103 | YES |
| 7X7E_Nb22 | 28GGT30 | -- | 102VPPGSRLRGC111 – V113 | YES |
| 7Z1A_H11 | 26GRTFSTA32 | 54SGGS57 | 100RVT102 – 104SL105 | YES |
| 7Z1B_A10 | 26GR27 – 30STA32 | 54SGGS57 | 100SAT102 – 104SLL106 | YES |
| 7Z1C_B5 | 25GRTFSTA31 | 53SGGS56 | 99QAT101 – 103SL104 – 110TWP112 | YES |
| 7Z1D_H11-H6 | 26GRT28 – 30STA32 | 54SGGS57 | 100KIT102 – 104SLL106 – 111TWP113 | YES |
| 7Z1E_H11-H4_mut | 25GRTFST30 | R51 – 53SGGS56 | 101VSYLLSD107 | YES |
| 8BEV_W25 | 26GSIFG30 | -- | 100KNELGF105 | YES |
| 8C8P_10D12 | 26GFT28 – L31 | 53SGG55 | 98GLGFGELPP106 | YES |
| 8CXN_Nb2-57 | -- | -- | -- | NO |
| 8CXQ_Nb1-22 | 26GRSFNS31 | 53GSPH56 | L102 – 104VGS106 | YES |
| 8CY6_Nb2-65 | 28GTIST32 | 55NLG57 | 102LEGGTQ107 | YES |
| 8CY7_Nb2-38 | 26AR27 – S29 | -- | 99GWGIRQP105 – I107 | YES |
| 8CY9_Nb1-23 | 26GRTDSI31 | 54SGGG57 | 100SLRVGS105 – S107 | YES |
| 8CYA_Nb2-67 | G26 | 53WNGSTR58 | 103DGVIDGTNANA113 | YES |
| 8CYB_Nb1-8 | 28GRTFSN33 | -- | 102RGSS105 | YES |
| 8CYC_Nb2-34 | 26GRTF29 | -- | 103YSRS106 | YES |
| 8CYD_Nb2-45 | 26GYD28 – 30SI31 | 52SRVGS56 | 99IPMTT103 | YES |
| 8CYJ_Nb1-25 | 26GR27 | 55VG56 | 103SSS105 | YES |
| 8CYJ_Nb2-10 | -- | -- | -- | NO |

| | | | | |
|---|---|---|---|---|
| 8CYJ_Nb2-62 | 26GR27 | 52ARS54 – 56DT57 | 101VIQYGIVPGND111 | YES |
| 8DI5_VHF6 | 24DF25 | -- | 98SGSG101 | YES |
| 8DLX_ab6 | G26 | -- | 99WLYGSGY105 | YES |
| 8DT8_LM18 | 25GFTF28 | 52GSGG55 | 102YGARDY107 | YES |
| 8DT8_Nb136 | 25GFT27 – 30SS31 | 52GSGGS56 | 100GPYDPTDSTY109 | YES |
| 8ELO_C4-225 | G26 | -- | 103YTYGGSV109 | YES |
| 8ELP_C4-240 | G26 | -- | 103YDQTGF108 | YES |
| 8ELQ_C4-255 | 26GF27 | 53GSGGS57 | 102YYSPYGGP109 | YES |
| 8EYG_NbUNK | 26GGTFSS31 | 54DGGA57 | 101VGKP104 | YES |
| 8G72_Nanosota2 | 26GFN28 – 31TS32 | 54GY55 – D57 | 99HNEPYFCDYSG109 | YES |
| 8G73_Nanosota3 | 26GSIFSP31 | -- | K100 | YES |
| 8G75_Nb4 | 26GF27 | 54SGGR57 | 102SRWYCPLQFSAD113 | YES |
| 8GZ5_VHH-P17 | 26GRTSS30 | 53GNGGT57 | -- | NO |
| 8H5T_Nb-015 | 25GFTLDS30 | 53DG54 | -- | NO |
| 8H5U_Nb-021 | 26GTGSTFST33 | -- | I102 | YES |
| 8H91_N19 | 25GGTFS29 | 51ADVGF55 | 98SLQSG102 | YES |
| 8HR2_Nb1B5 | 25GYTYST30 | -- | 98SGW100 | YES |
| 8HR2_Nb1C6 | 26GDTYSS31 | 54GGDN57 | 102CPWPDIGTMS111 | YES |
| 8K3K_Nb4 | 29GW30 – E32 | -- | -- | NO |
| 8OWT_A8 | 26GGT28 – 32TA32 | 53WRGVR57 | 100VGNYGL | YES |
| 8OWV_H6 | 26ESSLAP31 | D54 – 56HPTS59 | 108DS109 | YES |
| 8OWW_B5-5 | 26GSTL29 – N31 | 54RYGA57 | 101GPY103 | YES |
| 8Q7S_Ma6F06 | 25GITLDY30 | -- | 98GPLPPGHSCR107 – 109PT110 – 112LG113 | YES |
| 8Q7S_Re21H01 | 27GFT29 – S32 | 54ITGGS58 | 104RG105 | YES |
| 8Q93_Re21D01 | 26GFT28 - 30SSF32 | 52TITGGS57 | -- | NO |
| 8Q93_Re30H02 | 26GFTLDY31 | 53SSDSS57 | 100PATYYGGNWH109 | YES |
| 8Q94_Ma3B12 | 26GVT28 | E52 – 54SSGP57 | 104HEK106 – 110SPLG113 | YES |
| 8Q94_Re32D03 | 26GITLDY31 | -- | 99GPLPPGISC107 – T111 - 113LG114 | YES |
| 8Q95_Ma16B06 | 26GSI28 | 52PNSG55 | 99GVPV102 | YES |
| 8Q95_Ma3F05 | 26GVTLDG31 | 54SNGP57 | 104HER106 – S110 – 112LG113 | YES |
| 8RBY_Nb1.26 | 24GYT26 – 29SV30 | 50PSGRNR55 | 98SA99 – 101HDPE104 | YES |
| 8RJ7_Nb1.29 | 25GYTINT30 | 53GSG55 | 101GASG104 | YES |

| | | | | |
|---|---|---|---|---|
| 8SK5_VHH-7A9 | 26GGTAS30 | 54RNSGSTYV61 | 105PTLGWY110 | YES |
| 8ZER_P2C5 | 25GYTYC29 | -- | 99CSSGEYL105 | YES |

In this table, interacting patches predicted by our in-house method MLCE (REBELOT + BEPPE, https://github.com/colombolab/MLCE) are reported. For each of the 121 Nanobodies (Nbs) of our database, we report the PDB ID, the name given by the authors to the Nb, and the residues included in the predicted patches yielded at the end of the pipeline. For clarity, only residues belonging to CDR1, CDR2, or CDR3, predicted by state-of-the-art scheme Chothia using the Maestro Schrodinger suite (*see Material and Methods*), are reported, even though in most cases also framework residues were included in interacting patches. The only exceptions are 7FBJ_17F6 and 7FBK_20G6, for which Maestro could not find CDRs using the Chothia scheme, and for which we used the prediction of our NanoCDRx method as a reference.
Given the high variability of CDR3 sequence and structure, we were most interested in MLCE predictiveness for this loop. For this reason, we considered the prediction successful if at least one CDR3 residue was included in predicted patches. However, CDR1 and CDR2 predictions are also reported to completely assess MLCE performances.
Sequences of residues are indicated with the number of the first residue, followed by the one letter code of the residues, and then the number of the last residue (*e.g.* residues SLLS spanning from residue 104 to 107 are reported as 104SLLS107). The MLCE calculation is performed on the resolved 3D structure of Nbs: for this reason, the numbering of residues always starts from 1, with residue 1 being the first resolved residue in the crystal structure.

**Supplementary Figure 1 | Architecture of the nanocdr-x deep-learning model for nanobody sequence labelling**

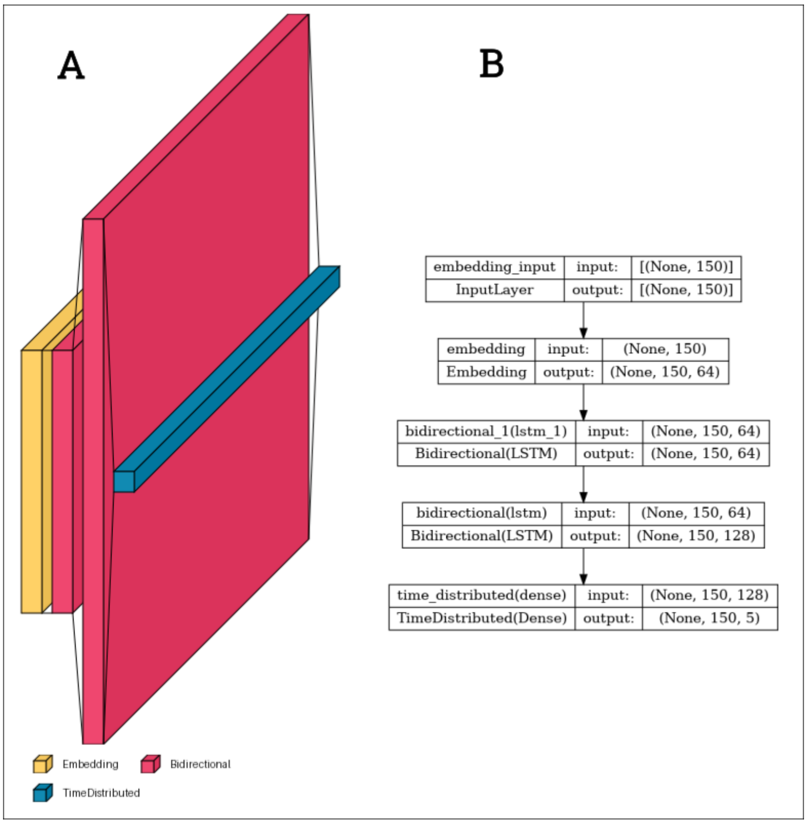


(A) Schematic 3D representation of the sequential architecture used for nanobody sequence labelling. The model is composed of an embedding layer (yellow), followed by two stacked bidirectional LSTM

layers (red), and a time-distributed dense output layer (blue). This configuration allows the network to learn contextual information from both N- and C-terminal directions of the sequence.
(B) Computational graph of the implemented architecture, showing the dimensionality and output shape at each layer. The embedding layer maps input tokens (sequence length 150) into a 64-dimensional dense space. The two BiLSTM layers, with 32 and 64 units respectively, process the sequence bidirectionally and return full sequences to preserve contextual dependencies. The final time-distributed dense layer applies a softmax activation to predict residue class probabilities across five categorical outputs (including padding).

**Supplementary Figure 2 | Training and validation performance of the nanocdr-x model across epochs**

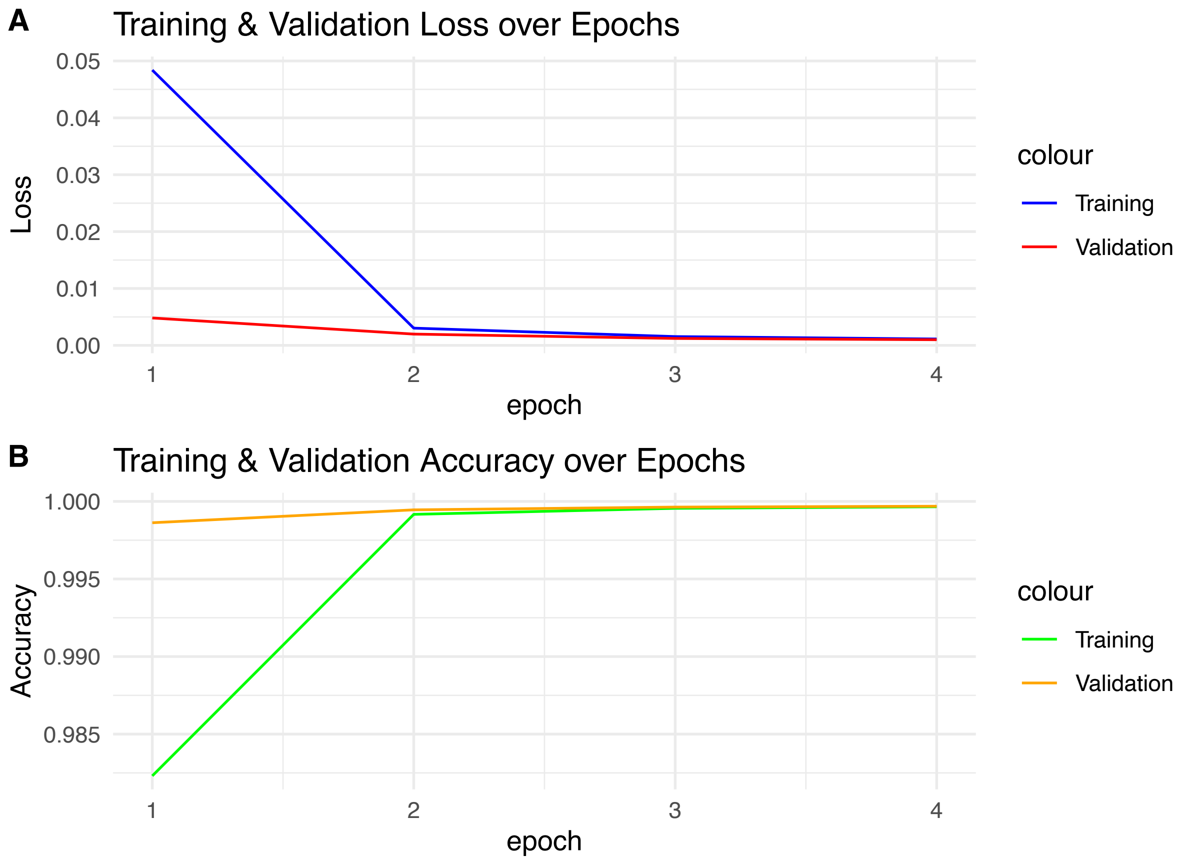


(A) Training and validation loss over four epochs. Both curves converge rapidly, with minimal loss reduction after epoch 2.
(B) Training and validation accuracy across epochs. Validation accuracy remains consistently higher than training accuracy, indicating the absence of overfitting. The model achieved a final validation accuracy of **99.9%** on the INDI dataset and was stopped after 4 epochs.
